# Lesions initiating spontaneous mitotic crossover are minimally subject to non-homologous end joining

**DOI:** 10.64898/2026.05.11.724412

**Authors:** Peter Chovanec, Shiyang He, Yi Yin

## Abstract

Homologous recombination (HR) between sister chromatids is the dominant outcome of replication-associated DNA repair, yet the lesions that initiate spontaneous mitotic crossovers remain poorly defined. Most mechanistic work on HR pathway choice uses enzymatically induced two-ended double-strand breaks (DSBs), where non-homologous end joining (NHEJ) is a major competing pathway. Whether NHEJ also competes for spontaneous, replication-associated lesions has not been directly tested. Here we use sci-L3-Strand-seq, a single-cell replication-template strand-specific sequencing method that maps sister chromatid exchange (SCE) genome-wide, to profile heterozygous and homozygous knockouts of NHEJ factors (LIG4, XRCC4) and the single-strand break (SSB) repair scaffold XRCC1 in HAP1 and BJ cell lines. NHEJ disruption produced only a modest (30%) increase in SCE, whereas XRCC1 loss caused a pronounced, 5-fold elevation. The dense, widespread SCE pattern in XRCC1-deficient cells is consistent with unrepaired SSBs being converted at replication forks into one-ended DSBs that lack a second end for ligation and therefore cannot engage NHEJ. In parallel, structural variation (SV) mapping in the same single cells revealed a strongly non-random landscape dominated by recurrent chromosome losses and clonal expansion, indicating selective pressure and stepwise genome evolution. SCE frequency did not correlate with SV burden or SV-defined subclones, demonstrating that error-free recombination and mutational rearrangement represent separable axes of genome maintenance. Recovery of reciprocal daughter-cell pairs with matching SCE breakpoints directly confirms that these events arose by inter-sister exchange in the preceding division. Together, these results show that spontaneous mitotic crossovers are driven by lesions largely incompatible with NHEJ and instead engage HR through replication-coupled SSB-to-DSB conversion, and that elevated error-free recombination is decoupled from the mutational SV landscape.

## Introduction

Large-scale structural variations (SVs), including copy-neutral loss-of-heterozygosity (cnLOH), copy-number variations (CNVs), inversions, and translocations are among the most consequential forms of genome alterations in cancer, hereditary disease, and evolution (Collins and Talkowski, 2025; Stankiewicz and Lupski, 2010), and have served as the principal phenotype through which spontaneous DNA repair fidelity is read out. Yet SVs are a retrospective and selective record: they capture only the rare repair events that misused an allelic inter-homolog or a non-allelic template, left a sequence-detectable scar, and survived clonal selection. A useful complementary lens is the topology of the joined product itself. Repair can either restore local continuity by rejoining one or both broken strands to the originally adjacent sequence or create non-local connectivity by re-establishing DNA continuity using a partner that was not contiguous along the same chromatid. Local-continuity outcomes include: 1) single-strand gap filling and nick ligation in base excision repair (BER), nucleotide excision repair (NER), and mismatch repair (MMR); 2) end-joining, including NHEJ and microhomology-mediated end joining (MMEJ); and 3) non-crossover gene conversion in HR. Non-local outcomes include SCE by HR between newly replicated sisters, end joining across isochromatid DSBs (simultaneous DSBs at the same locus on both sisters, which arise spontaneously from replication of a G1-phase DSB, as genetically observed in genome-wide cnLOH mapping in yeast (Lee et al., 2009; St Charles et al., 2012; Yin and Petes, 2013) and in immunoglobulin class switch recombination (Stavnezer et al., 2008), or from failed resolution of recombination intermediates (Yamamoto et al., 2011), and can be induced by Cas9 cleavage of both post-replicative sisters or by high-dose ionizing radiation), and long-range intra-chromatid deletions or inversion, all of which produce strand switches detectable by a single-cell template-strand sequencing method we and others have developed in recent years (Chovanec et al., 2026; Falconer et al., 2012; Hanlon et al., 2022). We operationally define SCE as a reciprocal strand switch between sister chromatids, without commitment to any particular repair mechanism. In **Fig.1A**, we depict SCE by HR and a second, mechanistically distinct route: double NHEJ (dNHEJ), in which end joining across an isochromatid DSB links the proximal end of one broken sister to the distal end of the other, producing a strand switch that mimics HR-mediated crossover. Apparent SCE outcomes, regardless of whether they arise from HR or NHEJ, are the primary focus of this study. SCE is by far the most frequent non-local outcome, because sister chromatids are the closest non-local partners, are held in proximity by cohesin from S phase through anaphase, and are sequence-identical; exchanges between them are therefore genetically silent and inconsequential. In mammalian cells, more than 99% (Johnson and Jasin, 2000) of crossover-type HR occurs between identical sisters and leaves no sequence change; the SVs visible to whole-genome sequencing (WGS) represent the remaining fewer than 1%, the detectable tip of a much larger non-local repair landscape. Sci-L3-Strand-seq is a high-throughput method that overcomes this limitation: by ablating BrdU-labeled nascent DNA strands and amplifying only parental template strands, it directly detects strand switches genome-wide and at scale, providing a readout of non-local DNA repair independent of any sequence change (Chovanec et al., 2026; Chovanec and Yin, 2025).

**Figure 1.**
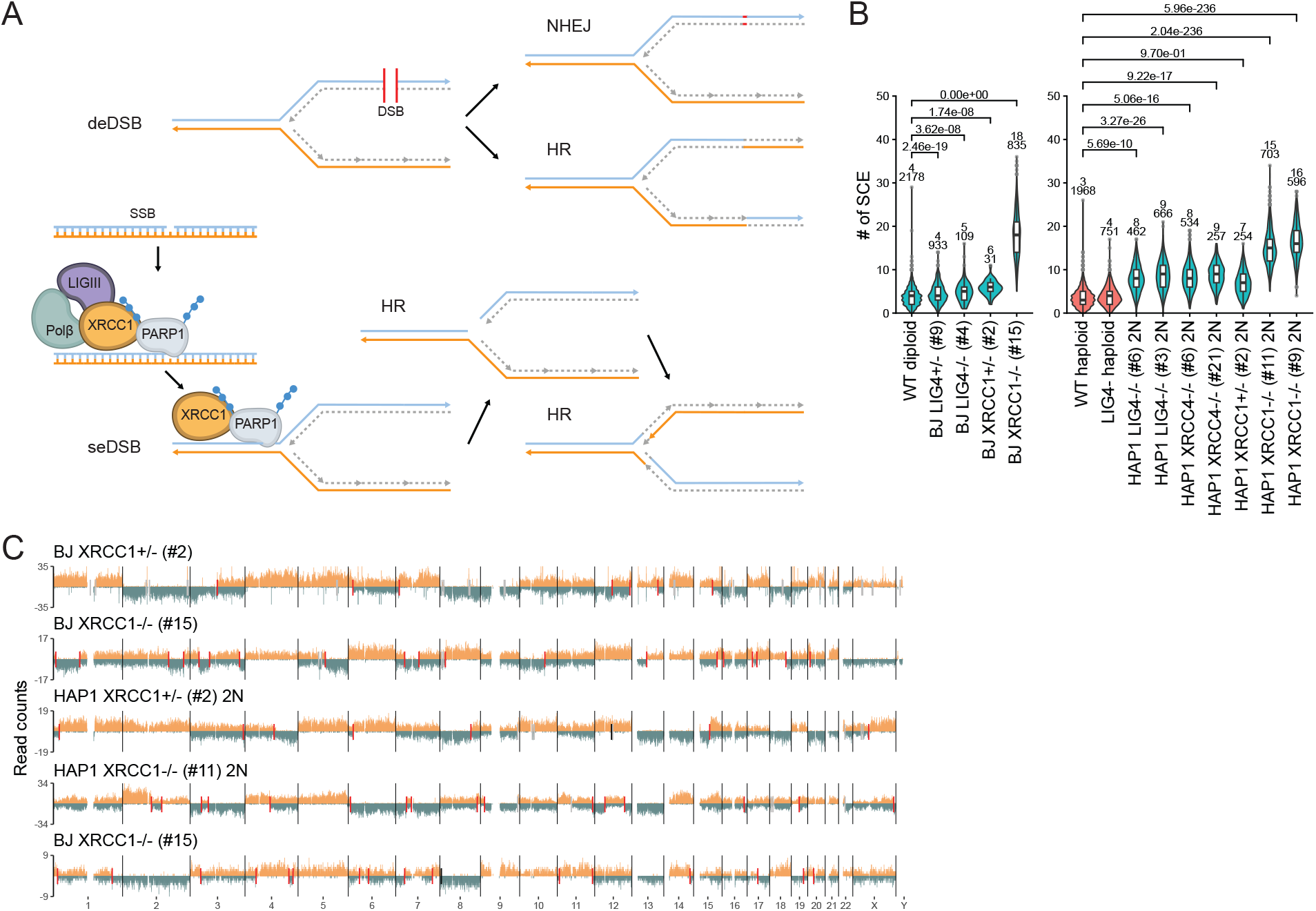
SCE increases in DNA repair–deficient cells. **A**. Single-ended (se) and double-ended (de) DSB in inducing SCE. **B**. Distribution of SCE counts per cell across wild-type and knockout lines in haploid (HAP1) and diploid (HAP1 2N and BJ-5ta) backgrounds. Significant difference was calculated using Mann-Whitney U with BH correction (p-values shown in plot). For comparisons with haploid cells, the number of SCEs were first normalized by ploidy. Numbers above each violin show the median number of SCEs and total number of cells. **C**. Representative genome-wide Strand-seq profiles from BJ-5ta wild-type, XRCC1+/-, and XRCC1-/- cells. SCEs are highlighted by red markers. Reads were counted across 200 Kb bins.

Single-strand breaks are among the most abundant endogenous DNA lesions, arising 10,000s of times per cell per day from oxidative damage, abortive topoisomerase I activity, and intermediates of BER (Caldecott, 2022). Their repair depends on XRCC1, which coordinates lesion detection and nick ligation through PARP1 and DNA ligase III (LIG3) (Caldecott, 2008; Demin et al., 2021). When this pathway fails, unligated SSBs persist into S phase and are converted by replication fork run-off into one-ended DSBs: structures that inherently lack a second end and are therefore poor substrates for classical NHEJ, which requires synapsis of two DNA termini by the Ku70/80–DNA-PKcs–XRCC4–LIG4 complex (Ciccia and Elledge, 2010; Zha and Yi, 2026). This structural asymmetry between replication-born, one-ended breaks and enzymatically-induced two-ended breaks (e.g., Cas9) predicts different pathway-choice outcomes. If the lesions that initiate spontaneous SCE are primarily two-ended DSBs, then loss of NHEJ should substantially increase SCE frequency by redirecting these breaks into HR, as has been observed at nuclease-induced DSBs (Chu et al., 2015). If instead a fraction of apparent SCEs reflects dNHEJ across two coincident, isochromatid DSBs (Waisertreiger et al., 2020), then NHEJ gene disruption should decrease the observed SCE frequency. Finally, if spontaneous SCE originates predominantly from one-ended, replication-born intermediates that cannot engage end joining, then NHEJ disruption should have minimal effect on SCE levels. Distinguishing among these alternatives requires comparing the SCE consequences of eliminating end joining (LIG4, XRCC4) with those of eliminating SSB repair (XRCC1) in otherwise isogenic backgrounds, while simultaneously monitoring structural variation to determine whether error-free recombination and mutagenic rearrangement are coupled or independent processes.

Here, we use sci-L3-Strand-seq to systematically compare how loss of NHEJ and SSB repair reshapes the non-local repair landscape at single-cell resolution. Using CRISPR knockouts and heterozygous lines in HAP1 and BJ cells, we show that disruption of NHEJ core factors LIG4 and XRCC4 produces only a modest increase in SCE, whereas loss of the SSB repair scaffold XRCC1 elevates SCE approximately five-fold. Integrated structural variation and clonal analysis in the same single cells reveals that SCE frequency and SV burden are uncorrelated, establishing error-free recombination and mutagenic rearrangement as separable axes of genome maintenance. Recovery of reciprocal daughter-cell pairs with matched strand-switch breakpoints directly assesses the origin of SCE events. Together, these results demonstrate that spontaneous mitotic crossovers are driven predominantly by replication-associated, NHEJ-incompatible lesions, and that the resulting recombination activity is decoupled from the structural variation landscape.

## Results

We generated CRISPR knockout lines targeting NHEJ factors including LIG4 and XRCC4 and the BER scaffold protein XRCC1, in both haploid (HAP1) and diploid (HAP1 2N and BJ) genetic backgrounds. As a benchmark, we included LIG4-deficient cells, for which we have previously demonstrated a modest but reproducible increase in SCE frequency using sci-L3-Strand-seq (Chovanec et al., 2026). This panel enabled a direct comparison of how disruption of mechanistically distinct repair pathways influences HR-mediated crossover events at single-cell resolution. Target loci were amplified and sequenced to validate CRISPR editing outcomes of individual clones (Supplemental Figures FigS1).

We used sci-L3-Strand-seq to measure how disruption of distinct DNA repair pathways alters SCE frequency (**Fig. 1B**). Wild-type cells exhibited low baseline levels: a median of 3 SCE events per cell in haploid HAP1 and 4 in diploid BJ-5ta (Chovanec et al., 2026). HAP1 cells frequently undergo spontaneous diploidization in culture, and the diploidized subpopulation showed a baseline of 6 SCE per cell, double the haploid value and consistent with the doubled chromosome content. Comparing the two diploids, diploidized HAP1 (6 SCE) had a higher baseline than BJ-5ta (4 SCE); as we have noted previously, we attribute this difference to their distinct cellular origins: HAP1 is a near-haploid derivative of the chronic myeloid leukemia line KBM-7, whereas BJ-5ta is a foreskin fibroblast line known for its exceptional karyotypic stability.

Loss of NHEJ genes resulted in a modest but consistent increase in SCE. LIG4^-/-^ haploid cells reached a median of 4 events, a 33% increase. Diploidized HAP1 LIG4^-/-^ and XRCC4^-/-^ clones reached medians of 8 to 9 events, 33-50% above the diploidized baseline. BJ-5ta LIG4^-/-^ increased from 4 to 5 (25%). In contrast, XRCC1 knockouts produced the highest SCE burden, with medians of 15 to 18 events per cell in HAP1 and BJ-5ta, corresponding to a 2.5- and 4.5-fold increase. Classical BrdU differential staining requires two successive rounds of replication and therefore captures SCE from two cell divisions. Our observed 15-18 SCE per single division is consistent with the ∼40 SCE previously reported by classical cytogenetics in XRCC1^-/-^ cells (Demin et al., 2021). We therefore conclude that one-ended, replication-born DSBs arising from unrepaired single-strand nicks (and/or gaps as a result of nuclease activity with the unligated nicks) are far more potent inducers of SCE (Pavani et al., 2024; Scully et al., 2024), and that NHEJ mainly competes with HR at two-ended DSBs, where its loss moderately increases SCE. We note, however, that a minor contribution of NHEJ to SCE cannot be excluded. The modest net increase observed in NHEJ-deficient cells may reflect the combination of a small reduction in SCE generated by dNHEJ between isochromatid DSBs and a larger increase due to redirection of two-ended breaks toward HR.

Interestingly, partial loss of XRCC1 in the heterozygous lines slightly increases SCE whereas heterozygous loss of LIG4 has no detectable effect on SCE frequency, indicating that SSB repair capacity is dosage-sensitive while NHEJ is not. These effects were consistent across independent knockout clones and cell lines, indicating that the observed differences reflect robust, pathway-specific modulation of recombination rather than experimental variation. We show representative example genome-wide single-cell profiles in **Fig.1C, FigS2** and **FigS3**.

We next annotated structural variants (SVs) for the BJ-5ta XRCC1^-/-^ (#15) line and grouped single cells into clusters based on sub-clonality of the SV events (**Fig2A**). The distribution of SVs across the genome was highly non-random, with a pronounced enrichment of copy number losses affecting chromosomes 4, 9, and 14. In addition, we identified a translocation in a subset of cells linking the terminal region of chromosome 9 to the proximal region of chromosome 14 (**Fig2B**). Notably, the clonal structure revealed cells carrying all combinations of losses across chromosomes 4, 9, and 14, including pairwise combinations (chr4/9, chr4/14, chr9/14) as well as cells harboring all three events. This pattern is consistent with a gradual, stepwise accumulation of SVs during clonal expansion rather than the emergence of a single catastrophic event. Together, these observations indicate that SVs arise non-randomly, with specific chromosomes showing recurrent events across cells. Importantly, these patterns do not correspond to the genomic location of the XRCC1 locus on chromosome 19, suggesting that they are unlikely to be a direct consequence of the editing target itself. Whether these recurrent losses reflect intrinsic chromosomal fragility, selective advantage conferred by reduced gene dosage, or both, cannot be distinguished from a single time-point sampling and remains an open question.

**Figure 2.**
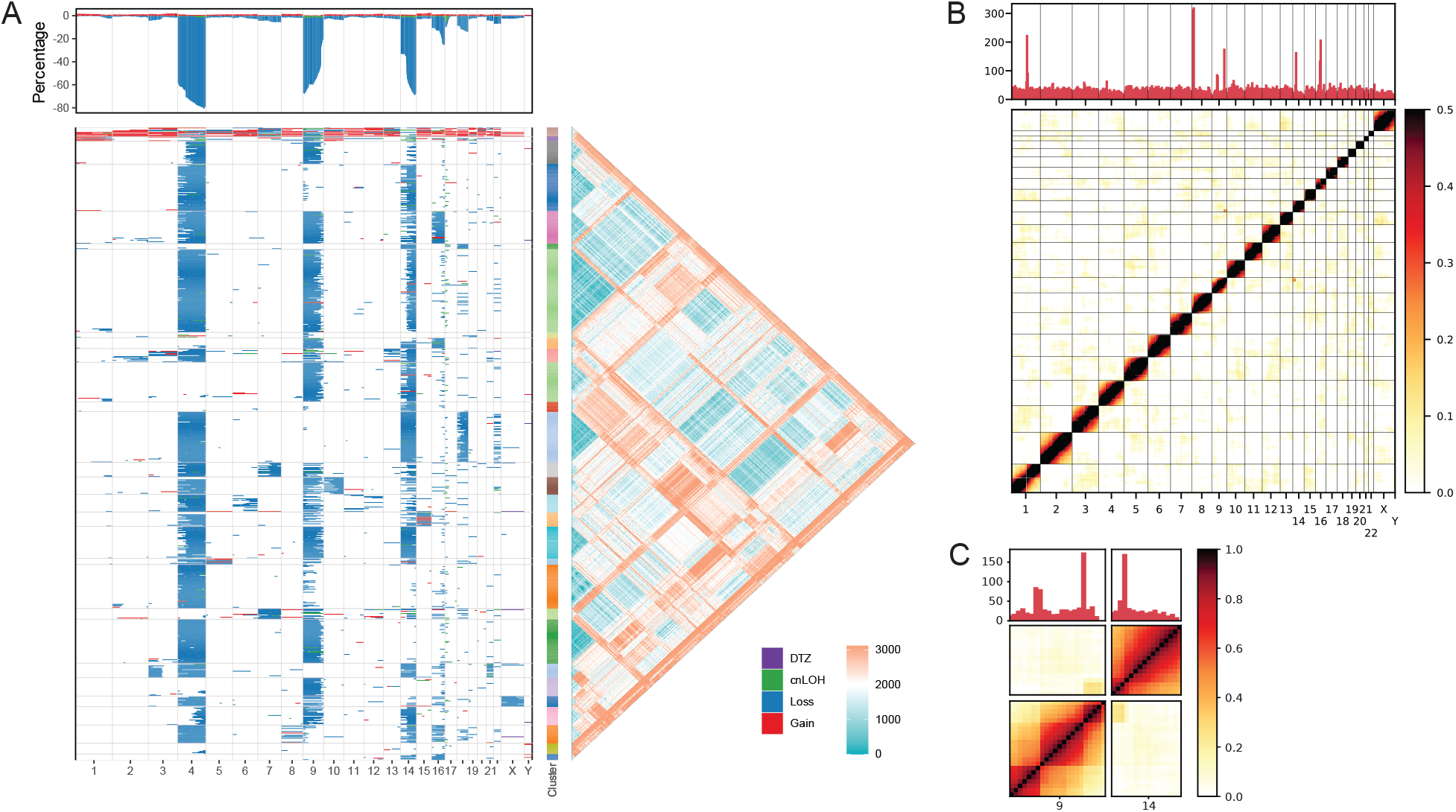
Structural variations in the XRCC1 knockout. **A**. Heatmap of SV annotations for the BJ-5ta XRCC1^−^/^−^(#15) line. Only cells with at least one SV annotation are shown (832/835). Each row represents individual cells grouped into clusters based on SV profiles using hierarchical clustering. Each cluster is uniquely colored to the right, along with the Canberra distance matrix used for clustering on the left, highlighting similarity between individual clusters. **B**. Genome-wide heatmap of strand-state Pearson positive correlation in BJ-5ta XRCC1^−^/^−^(#15) cells. Heatmap was plotted at a 5 Mb bin resolution using all 835 cells. Strand switches without the region filter are plotted in the top panel. Positive correlation in trans (off-diagonal) highlights mapped translocations. **C**. SCE frequency across SV-defined clusters. Boxplots show median SCE counts per cluster stratified by SV composition, demonstrating no significant association between SCE burden and SV state.

Despite this structured SV landscape, we found that SCE frequency did not correlate with any specific combination of SVs (**Fig2C**) unlike previously observed for a mouse fibroblast Patski cell line (Chovanec et al., 2026). Clusters lacking detectable SVs exhibited comparable SCE levels to those containing one or multiple SV events, indicating that the presence or accumulation of large-scale structural alterations does not measurably influence the observed single cell division SCE rates in this context. These findings suggest that, in XRCC1-deficient cells, SCEs arise largely independently of pre-existing SVs, highlighting a context-dependent relationship between recombination and genome instability. In certain cell lines like Patski, SCE and SV can represent positively correlated fragility for certain regions of the genome, whereas here SCE forms an orthogonal dimension from SV accumulation.

Finally, if SCE arises through reciprocal mitotic crossover, the resulting products should be reciprocal and copy-neutral, lacking the large-scale deletions or amplifications characteristic of catastrophic genome instability events driven by randomly joined DNA ends (Umbreit et al., 2020; Zhang et al., 2015) or by a half-crossover mechanism defined in yeast genetics, where one daughter cell gets a linkage switch (like SCE/strand-switch) while the other gets a terminal deletion from the switch point to the chromosome end, characteristic of defective break-induced replication, as observed in pol32 mutants in yeast (Deem et al., 2008). Reciprocal daughter cell pairs (RDCP) have been analyzed to capture all the four post-replication chromatids in HR for over 20 years in yeast (Barbera and Petes, 2006; St Charles et al., 2012; Yin et al., 2017) and for studying SVs in mammalian cells (Corazzi et al., 2026; Cosenza et al., 2025; Umbreit et al., 2020; Zhang et al., 2015). Because sci-L3-Strand-seq is high-throughput in nature, we reasoned that RDCPs could be identified directly when a significant proportion of daughter cells formed in one division are sampled. In **Fig.3A** and **FigS4**, we show three RDCPs that together harbor reciprocal products of 87 spontaneous SCE events in the XRCC1^-/-^. In contrast, in a companion paper (Chovanec *et. al*., 2026, manuscript submitted) where we analyzed Cas9-induced SCE events with sgRNA targeting repetitive sites in the genome, RDCPs frequently exhibited non-reciprocal SVs, as a result of the repair of two-ended DSBs. We therefore conclude that spontaneous SCEs manifested in the XRCC1^-/-^ background are not only more frequent but also qualitatively distinct from SCE-like events induced by Cas9-generated two-ended DSBs.

**Figure 3.**
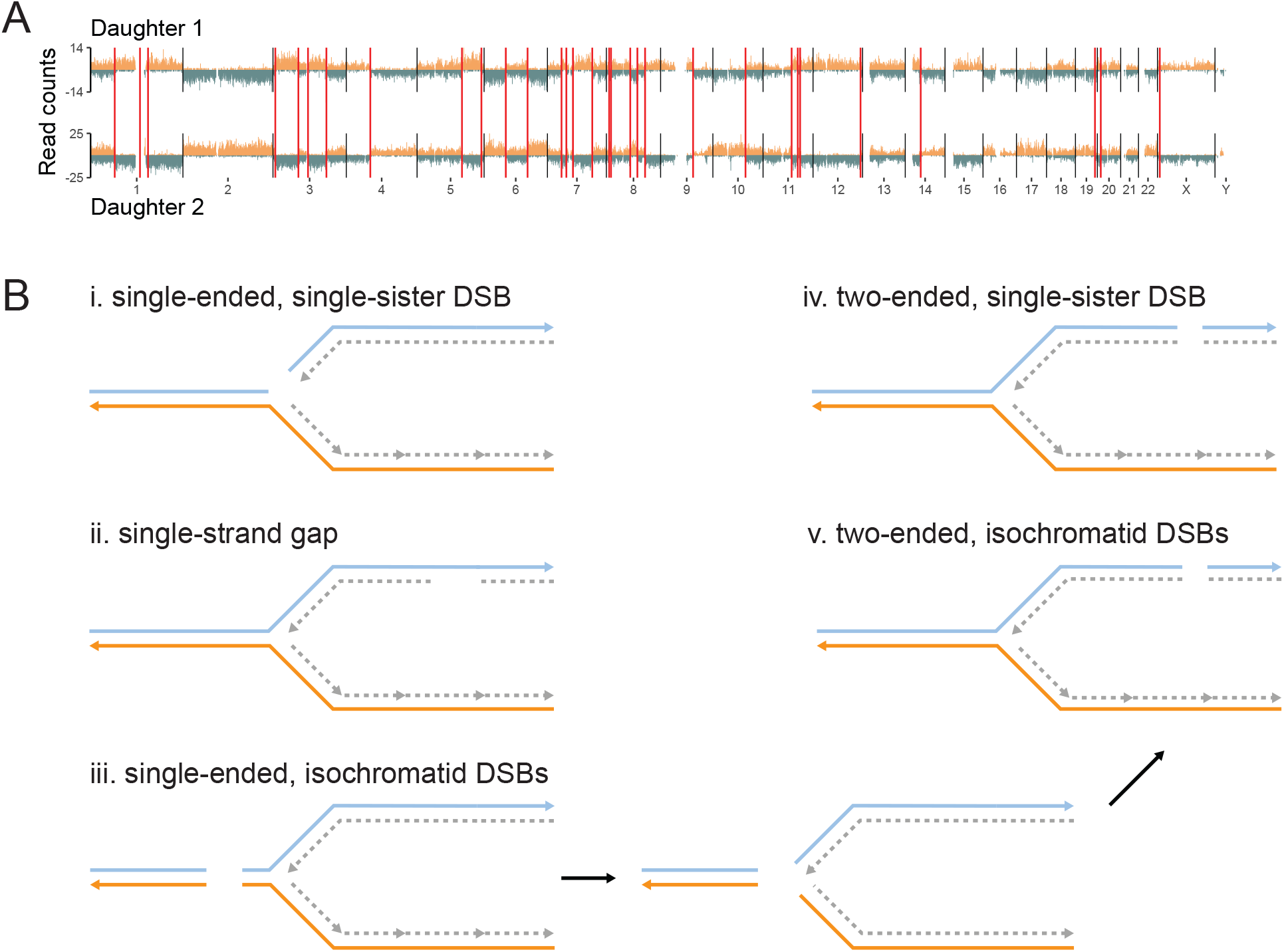
Reciprocal daughter cell pair (RDCP) analysis. **A**. Capture of reciprocal daughter cells. Each vertical red line highlights a reciprocal SCE event. This example RDCP has 30 SCEs. **B**. Five Classes of DNA substrate that can potentially generate SCE outcomes.

## Discussion

Non-local DNA repair outcomes, defined by the re-establishment of continuity using a partner that was not originally contiguous along the same chromatid, produce strand switches detectable by template-strand sequencing. Among these, SCE is by far the most frequent: it is reciprocal, copy-neutral, and genetically silent, representing a benign resolution that leaves no mutational footprint. This study set out to determine what lesion type initiates this predominant non-local outcome and whether NHEJ competes for the same substrates. By comparing NHEJ-deficient and SSB repair-deficient cells, we show that: 1) XRCC1 loss elevates SCE approximately five-fold, establishing unrepaired SSBs converted to one-ended, replication-born DSBs as the dominant initiating lesion; 2) NHEJ disruption increases SCE by only 25-50%, and critically does not decrease it, demonstrating that spontaneous SCE is generated by HR rather than by dNHEJ across isochromatid breaks; 3) this modest NHEJ-dependent increase is highly reproducible across independent clones, cell lines, and ploidy backgrounds, confirming that sci-L3-Strand-seq, leveraging its throughput, resolves subtle, pathway-specific differences; 4) SCE frequency does not correlate with SV, establishing these as orthogonal dimensions of genome maintenance; and 5) RDCP analysis recovered both daughter cell products harboring 87 SCE events. They are uniformly reciprocal, copy-neutral, and free of associated large SVs, directly confirming their origin as error-free inter-sister crossovers.

### DNA substrate that can potentially generate SCE

In **Fig.3B**, we conceptualize five classes of DNA substrate that could generate SCE, defined here as a genomic outcome (strand switch between sisters) regardless of mechanism. These substrates differ in two key properties: single-ended versus two-ended, and whether one or both sisters are affected. For single-ended substrates: 1) a replication-born DSB on one sister, caused by fork run-off at an unrepaired single-strand nick; 2) a single-stranded gap from incomplete replication or nuclease expansion of a nick; and 3) single-ended isochromatid DSBs, in which both sisters are broken at approximately the same position. For two-ended substrates: 4) a two-ended DSB on one sister, the classical competing substrate between NHEJ and HR; and 5) a two-ended isochromatid DSB, as produced by Cas9 cleavage of both post-replicative sisters, which could generate apparent SCE by dNHEJ. Below we consider each in turn.

Class i and iv: single-ended vs. two-ended DSB. Our comparison of XRCC1 and NHEJ knockouts directly supports that the former is the major source of spontaneous SCE. Loss of XRCC1 elevates SCE approximately five-fold, consistent with a large increase in spontaneous SSBs that persist into S phase to generate one-ended, replication-born DSBs. In addition, loss of XRCC1 promotes PARP1/2 trapping at nicks (Demin et al., 2021; Zhang and Zha, 2024), further increasing fork breakage. In contrast, if two-ended DSBs were the primary SCE-initiating lesion, NHEJ loss should redirect a large fraction of breaks into HR. Instead, NHEJ disruption increases SCE by only 25-50%, indicating that two-ended DSBs, the classical competing substrate between NHEJ and HR, make a limited contribution to spontaneous SCE. We note that this study specifically tests classical NHEJ; whether MMEJ competes with HR for one-ended replication-born DSBs is not addressed by our perturbations and remains an open question.

Class ii: single-strand gaps arising from incomplete replication (Ulrich and Jentsch, 2000) or from nuclease expansion of unligated nicks, as demonstrated for Exo1 processing of NER intermediates (Giannattasio et al., 2010) and MRN-mediated expansion of PARP inhibitor-trapped nicks (Cong et al., 2021; Seppa et al., 2025), represent a second potential SCE-potent substrate. Such gaps could generate SCE through UBC13/MMS2-dependent template switching (Daigaku et al., 2010), in which the nascent strand on the stalled fork invades the replicated sister to bypass the lesion. However, whether and how the resulting joint molecules are resolved as crossovers remains unclear. Our study does not directly test this substrate.

Class iii & v: single-ended isochromatid DSBs affecting both sister chromatids. Such breaks appear single ended: there is no pair of ends available for NHEJ on either sister. Whether this configuration produces SCE depends on the converging fork. If TRAIP-mediated unloading of the CMG helicase (Deng et al., 2019; Sonneville et al., 2019; Wu et al., 2021) allows nuclease processing of the converging fork terminus and the resulting ends are proximal enough, trans end joining between sisters could generate a strand switch. Alternatively, if converging fork converts the blocked fork into two two-ended DSBs, similar to Class v, because neither sister is intact, both may still require repair using the allelic homolog or non-allelic repeats, potentially causing cnLOH, CNVs, inversions or translocations. In budding yeast, G1 DSBs replicated into isochromatid breaks are repaired by inter-homolog HR, producing 4:0 gene conversion rather than the 3:1 pattern expected from a single broken chromatid (Lee et al., 2009; St Charles et al., 2012; Yin and Petes, 2013). The small yeast nucleus, Rabl chromosome organization (Duan et al., 2010), and somatic homolog pairing (Kim et al., 2017; Weiner and Kleckner, 1994) may make inter-homolog repair more accessible in yeast than in mammalian cells, where homologs are spatially separated. Consistent with this, Waisertreiger et al. (Waisertreiger et al., 2020) found that when both alleles at a fragile site are broken under replication stress, DSBs persist into mitosis with no available template. In human cells, the ATM-dependent G1/S checkpoint is stringent, and G1 DSBs are normally repaired or trigger arrest before S phase; a notable exception is class switch recombination, where ATM signaling is specifically dampened to permit AID-induced G1 breaks to persist into replication (Stavnezer et al., 2008). Outside this specialized context, replication of G1 breaks and the consequent formation of isochromatid DSBs is expected to be rare. Nevertheless, closely-opposed lesions (single-stranded in nature) ahead of the fork, as well as interstrand crosslinks by aldehyde metabolites, can break both sisters in S phase, causing apparent SCE by dNHEJ: end joining of the proximal end of one broken sister to the distal end of the other generates a strand switch indistinguishable from HR-mediated crossover. If dNHEJ were a significant contributor to spontaneous SCE, then NHEJ loss should decrease the observed SCE frequency. We observe the opposite: SCE increases modestly in LIG4^-/-^ and XRCC4^-/-^ cells, ruling out dNHEJ as a major contributor and establishing that spontaneous SCE is overwhelmingly generated by HR.

### SCE and SV represent fundamentally different measurements

SCEs detected by sci-L3-Strand-seq were formed in the immediately preceding cell division: it is a snapshot of current repair activity. SVs, by contrast, are the cumulative, heritable record of rare mutagenic events that have survived clonal selection over many generations. SCE is therefore a real-time readout of repair pathway usage, while the SV landscape is a historical record shaped by both mutation and selection. Our prior analysis of the Patski mouse fibroblasts revealed a positive correlation of SCE and SV at specific genomic loci (Chovanec et al., 2026), suggesting that in some contexts the same underlying fragility drives both or certain pre-existing SVs drive SCE in a chromosome-autonomous fashion. While Patski is an established cell line with accumulated and selected SVs, SVs in the XRCC1 knockout line are newly acquired upon engineered loss of BER. That these two dimensions are uncorrelated in XRCC1^-/-^ cells (SV-defined subclones show indistinguishable SCE levels regardless of their structural variant load) confirms that they reflect independent processes.

### Recovery of RDCPs reveals the reciprocal nature of SCE and a lack of large-scale SVs in the XRCC1^-/-^

The 87 SCE events captured across three XRCC1^-/-^ RDCPs are reciprocal and copy-neutral. Both daughter cells carry the expected complementary strand-state configurations, the hallmark of “healthy” crossover: the genome undergoes strand exchange without any net gain, loss, or rearrangement of sequence. The quality of these spontaneous crossovers stands in sharp contrast to three other settings. First, in a companion paper analyzing Cas9-induced events with sgRNA targeting repeats, RDCPs frequently exhibit non-reciprocal SVs. Second, in yeast BIR studies, *pol32* mutants produce half-crossovers: one daughter cell inherits a linkage switch (resembling an SCE strand-switch) while the other suffers a terminal deletion from the switch point to the chromosome end (Deem et al., 2008). In both cases, the lesion structure (two-ended or pathological BIR) permits large SVs. The clean reciprocity of XRCC1^-/-^ SCEs argues that the one-ended, replication-born DSBs driving these events are channeled exclusively through high-fidelity inter-sister HR.

A striking parallel exists in yeast. We previously mapped UV-induced mitotic recombination genome-wide and recovered ∼381 cnLOH events (300 gene conversions, 60 crossovers, 21 BIR), virtually all copy-neutral, with no large deletions, duplications, or translocations detected (Yin and Petes, 2013). The mechanism is analogous to what we observe: UV-induced pyrimidine dimers are processed by NER into nicks; when these nicks persist into S phase, replication fork run-off generates one-ended DSBs that are repaired by HR. Constrained by single-cell sequencing technologies, only inter-homolog HR was analyzed; yet perturbations of various polymerases, checkpoint genes, and topoisomerases and the two RNaseH genes are all accompanied by strong CNVs in addition to cnLOH (Andersen et al., 2015; Deng et al., 2015; McCulley and Petes, 2010; Quevedo et al., 2015; Song et al., 2014; Zhang et al., 2022; Zheng et al., 2016), indicating non-allelic repair and gross chromosomal rearrangements. In both yeast and human, the outcomes are overwhelmingly copy-neutral. The comparison illustrates a broader principle: the initiating lesion dictates the execution of repair, determining not only pathway choice (end joining vs. HR) but also partner choice (sister vs. homolog vs. non-allelic) and outcome fidelity. One-ended, replication-born breaks are intrinsically channeled toward high-fidelity recombination because their structure excludes end joining and their proximity to the sister chromatid favors the identical template.

### Limitations

Several limitations should be noted. Sci-L3-Strand-seq detects SCE at approximately 200 kb resolution; indels, non-crossover gene conversions, and closely spaced double crossovers would be invisible. The distinction between true SCE by HR and apparent SCE by dNHEJ relies on indirect inference from genetic perturbation; direct characterization of junction sequences is lacking.

## Conclusion

In summary, by comparing NHEJ-deficient and SSB repair-deficient cells using single-cell template-strand sequencing, we demonstrate that spontaneous mitotic crossovers are initiated predominantly by one-ended, replication-born DSBs that are structurally incompatible with NHEJ. The observed SCE is predominately generated by HR, not by dNHEJ across isochromatid breaks, and is overwhelmingly reciprocal and copy-neutral. SCE frequency is decoupled from the SV landscape, establishing error-free recombination and mutagenic rearrangement as independent axes of genome maintenance. These findings reframe pathway choice in spontaneous DNA repair: the dominant competition is not between HR and NHEJ for the same two-ended break, but between SSB repair (preventing the lesion) and HR (repairing the replication-born consequence), with NHEJ largely excluded by the one-ended structure of the initiating break. Just as UV-induced nicks in yeast produce clean, copy-neutral HR rather than CNVs, unrepaired SSBs in human cells generate reciprocal SCEs that leave no large-scale mutational footprint. The lesion shapes the execution.

## Acknowledgments

This work was supported by National Institute of General Medical Sciences (R35GM142511 to Y.Y) and W. M. Keck Foundation (1002444 to Y.Y).

## Author Contributions

Conceptualization: P.C., Y.Y.; Data curation: P.C, S.H., Y.Y.; Software: P.C., Y.Y.; Formal analysis: P.C., Y.Y.; Funding acquisition: Y.Y.; Investigation: P.C., S.H., Y.Y.; Methodology: P.C., S.H., Y.Y.; Supervision: Y.Y.; Writing - original draft: P.C., Y.Y.; Writing - review & editing: P.C., S.H., Y.Y.

## Methods

### Cell culture

BJ-5ta (CRL-4001, ATCC) cells were cultured in DMEM/F12 medium supplemented with 10% FBS and 1x Pen-Strep. HAP1 cells were cultured in IMDM medium supplemented with 10% FBS and 1x Pen-Strep. All cells were cultured at 37°C, 5% CO_2_.

### Generation of HAP1 and BJ-5ta knockout clones

Guide RNAs (gRNA) were designed using CHOPCHOP v3 (Labun et al., 2019). Knockout clones were generated using the Alt-R S.p. Cas9 Nuclease V3 (1081058; IDT) with Alt-R CRISPR-Cas9 tracrRNA (1072532; IDT) and Alt-R CRISPR-Cas9 crRNA (LIG4: 5’-GCTTATACGGATGATCATAA-3’; XRCC4: 5’-ATGGTCATTCAGCATGGACT-3’; XRCC1: 5’-TGCAGGACACGACATGG-3’; LIG3: 5’-CTGTTAGGTACACATCACCG-3’) following the manufacturer’s protocol. Briefly, crRNA and tracrRNA were diluted to 1µM with duplex buffer (11010301; IDT) and annealed by heating to 95°C for 5 minutes and cooled to room temperature (RT) on benchtop for 30 minutes to form the gRNA. Alt-R S.p Cas9 enzyme was diluted to 1µM with OptiMEM, combined with gRNA and Cas9 PLUS reagent (CRISPRMAX kit, CMAX00003; Invitrogen), and incubated at RT for 5 minutes to form the RNP complex. Reverse transfection was performed with cells diluted to 200,000 cells/mL. Following a 48 hour incubation at 37°C 5% CO_2_, cells were single-cell sorted with FACS into 96 well plates. For HAP1, cells were gated on low FSC/SSC to select for haploid cells. After 15-30 days, individual clones were genotyped with either T7 Endonuclease I pre-screening (M0302; NEB) and Sanger sequencing or amplicon sequencing as previously described. Oligos used for genotyping are provided in TableS1. Sanger sequencing was analysed in Benchling. Amplicon sequencing was analyzed using CRISPResso2.

### Cell labeling and Sci-L3-Strand-seq library generation

All cells were cultured with BrdU at 40 µM final concentration for 24 hours prior to fixation. The sci-L3-Strand-seq protocol was carried out as previously described (Chovanec and Yin, 2025; Yin et al., 2019). Briefly, cells were trypsinized and fixed with 37% formaldehyde (1.5% final concentration) in 1x PBS at a cell density of 1 million/mL for 10 minutes at room temperature with gentle tube inversion, followed by quenching with 100 µL of 2.5 M glycine per 1 mL of FA. After fixation, cells were snap frozen with liquid nitrogen in nuclei freezing buffer (NFB: 50mM Tris (pH 8.0), 25% glycerol, 5mM Magnesium acetate Mg(OAc)2, 0.1mM EDTA, 5mM DTT, 1x Protease inhibitor cocktail (Sigma, P8340)). Tagmentation was performed with 24 first round barcodes using Tn5 transposase as previously described (Yin et al., 2019). First round barcode adapters were loaded by mixing 1 µL of diluted Tn5 with 1.4 µL of 1.5 µM annealed adapters, followed by incubation at room temperature for 30 minutes. After tagmentation at 55 °C for 15 minutes, the reaction was stopped with the addition of LBT (lysis buffer with triton: 60 mM Tris-Ac pH 8.3, 2 mM EDTA pH 8.0, 15 mM DTT, 0.1% Triton-X100), followed by nuclei pooling. The subsequent ligation was performed with 72 second round barcodes. Ligation was stopped by the addition of LBT and the pooled nuclei were stained with Hoechst-33258 to a final concentration of 10ng/µL. The quenched population was sorted with FACS (200-300 nuclei per well) into 96-well plates containing 3 µL of LBT. Sorted plates were stored in -80 °C. After gap extension, Hoechst-33258 was again added at a final concentration of 10ng/µL and incubated at room temperature for 10 minutes in the dark. A dose of 270 mJ/cm^2^ was administered with a UVP crosslinker (CL-3000L, 365 nm). After UV treatment, 0.9uL of diluted USER enzyme (M5505L, NEB; 1:3 dilution) was added and incubated at 37°C for 15 minutes, followed by the addition of 0.9 µL of Uracil Glycosylase Inhibitor (UGI) (M0281; NEB) and a further incubation at 37 °C for 10 minutes. Finally, the T7 in vitro transcription system was assembled by adding 2 µL H_2_O, 2 µL T7 Pol mix and 10 µL rNMP mix (E2050; NEB, HiScribe T7 Quick High Yield RNA Synthesis Kit) and incubated at 37°C for 14-16 hours. After RNA purification, reverse transcription, and second strand synthesis (SSS) that incorporates the third round barcode, sequencing libraries were prepared using the NEBNext Ultra II DNA Library Prep Kit for Illumina (E7645S, NEB) with NEBNext Multiplex Oligos for Illumina (Index Primers Set 2) (E7500S, NEB) following the manufacturers protocol.

### Breakpoint segmentation, SCE and de novo SVs annotation for sci-L3-Strand-seq

Raw fastq’s were processed using the sciL3Pipe snakemake pipeline (v0.2.1) (https://github.com/recombinationlab/sciL3Pipe), which de-multiplexes the split and pool barcodes, performs genome alignment, filtering, and produces individual BAMs for each single cell. Single-cell BAMs were subject to breakpoint identification and filtering using breakpointR (v1.10.0) and sciStrandR (v0.2.3). Cells with passing quality metrics (background estimate > 0 and <= 0.08, strand-neutral < 0.75, median reads per Mb > 2) were used in downstream analysis. Translocations and inversions were identified as previously described, with the exception of all WW and CC regions being used for inversions, instead of the previous exclusion of chromosomes with breakpoints. New breakpoint hotspots in one or more clones matching inversions or translocations were added to the region filter. Segmentation from strand-state breakpoints were used for SV annotation using a support vector machines (SVM) classifier with strand state, coverage, digital counts, and haplotype as the input features.

### Reciprocal daughter cell identification

The genome is tiled into 1 Mb bins across autosomes and chrX. For each QC-passing cell, W and C reads are counted per bin, and a strand ratio score is computed as (W − C)/(W + C), yielding +1 for pure W and −1 for pure C. Bins with fewer than 4 reads are set to 0. For each pair of cells, a sisterness score is calculated: at bins where at least one cell has |score| > 0.5 (i.e., a confident non-WC state), the score is |score_i + score_j|. This sum approaches 0 when the two cells have opposite strand states (one WW, one CC — the expected reciprocal pattern for sister cells) and approaches 2 when they share the same state. The sisterness score is the mean of this quantity across all informative bins; lower values indicate stronger sister cell candidates. The top 20 pairs with the lowest sisterness scores are reported, and strand-seq ideograms for each pair are plotted side-by-side for visual confirmation of the reciprocal strand inheritance pattern.

