## Supplemental Figures for "Lesions initiating spontaneous mitotic crossover are minimally subject to non-homologous end joining"

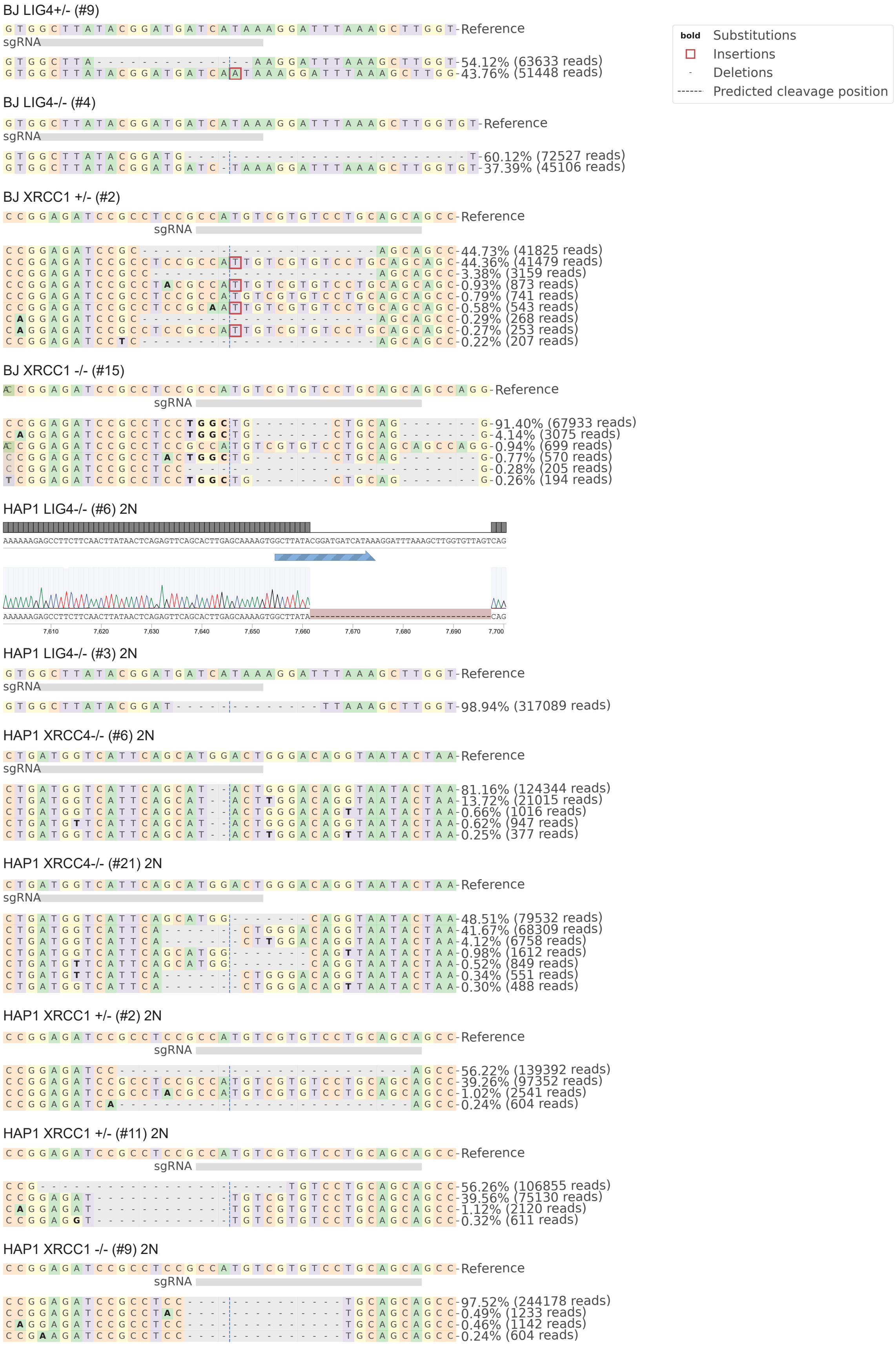

BJ LIG4+/- (#9)
Reference
sgRNA
54.12% (63633 reads)
43.76% (51448 reads)
bold Substitutions
Insertions
Deletions
Predicted cleavage position
BJ LIG4-/- (#4)
Reference
sgRNA
60.12% (72527 reads)
37.39% (45106 reads)
BJ XRCC1 +/- (#2)
Reference
sgRNA
44.73% (41825 reads)
44.36% (41479 reads)
3.38% (3159 reads)
0.93% (873 reads)
0.79% (741 reads)
0.58% (543 reads)
0.29% (268 reads)
0.27% (253 reads)
0.22% (207 reads)
BJ XRCC1 -/- (#15)
Reference
sgRNA
91.40% (67933 reads)
4.14% (3075 reads)
0.94% (699 reads)
0.77% (570 reads)
0.28% (205 reads)
0.26% (194 reads)
HAP1 LIG4-/- (#6) 2N
AAAAAAGAGCCTTCTTCAACTTATAACTCAGAGTTCAGCACTTGAGCAAAAGTGGCTTATACGGATGATCATAAAGGATTTAAAGCTTGGTGTTAGTCAG
AAAAAAGAGCCTTCTTCAACTTATAACTCAGAGTTCAGCACTTGAGCAAAAGTGGCTTATA
CAG
7,610
7,620
7,630
7,640
7,650
7,660
7,670
7,680
7,690
7,700
HAP1 LIG4-/- (#3) 2N
Reference
sgRNA
98.94% (317089 reads)
HAP1 XRCC4-/- (#6) 2N
Reference
sgRNA
81.16% (124344 reads)
13.72% (21015 reads)
0.66% (1016 reads)
0.62% (947 reads)
0.25% (377 reads)
HAP1 XRCC4-/- (#21) 2N
Reference
sgRNA
48.51% (79532 reads)
41.67% (68309 reads)
4.12% (6758 reads)
0.98% (1612 reads)
0.52% (849 reads)
0.34% (551 reads)
0.30% (488 reads)
HAP1 XRCC1 +/- (#2) 2N
Reference
sgRNA
56.22% (139392 reads)
39.26% (97352 reads)
1.02% (2541 reads)
0.24% (604 reads)
HAP1 XRCC1 +/- (#11) 2N
Reference
sgRNA
56.26% (106855 reads)
39.56% (75130 reads)
1.12% (2120 reads)
0.32% (611 reads)
HAP1 XRCC1 -/- (#9) 2N
Reference
sgRNA
97.52% (244178 reads)
0.49% (1233 reads)
0.46% (1142 reads)
0.24% (604 reads)

**Figure S1**. Genotyping of CRISPR-Cas9 edited clones with amplicon sequencing. Target loci were amplified and sequenced to validate CRISPR–Cas9 editing outcomes of individual clones. Allele frequency tables around each sgRNA generated by CRISPResso2 and a Sanger sequencing trace (Benchling) highlight the edits present within each clonal line.

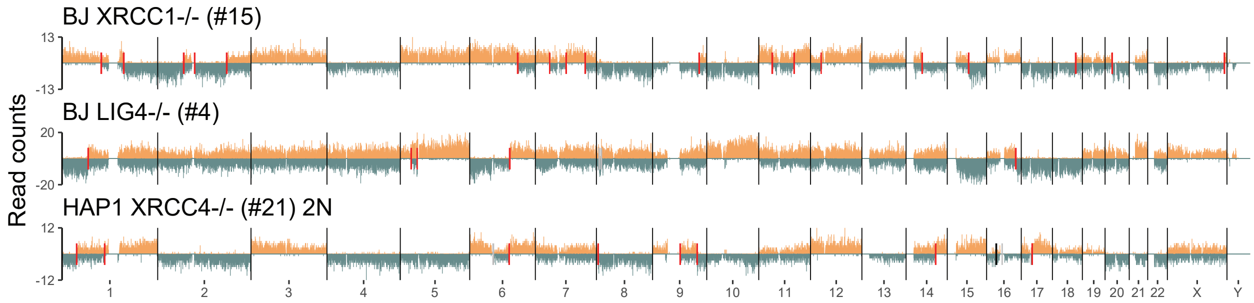


**Figure S2.** Additional examples of single-cell Strand-seq profiles across DNA repair knockouts. Representative genome-wide Strand-seq profiles from independent clones of LIG4$^{-/-}$, XRCC4$^{-/-}$, XRCC1$^{+/-}$, and XRCC1$^{-/-}$ cells in both HAP1 and BJ-5ta backgrounds. SCEs are highlighted by red markers. Reads were counted across 200 Kb bins.

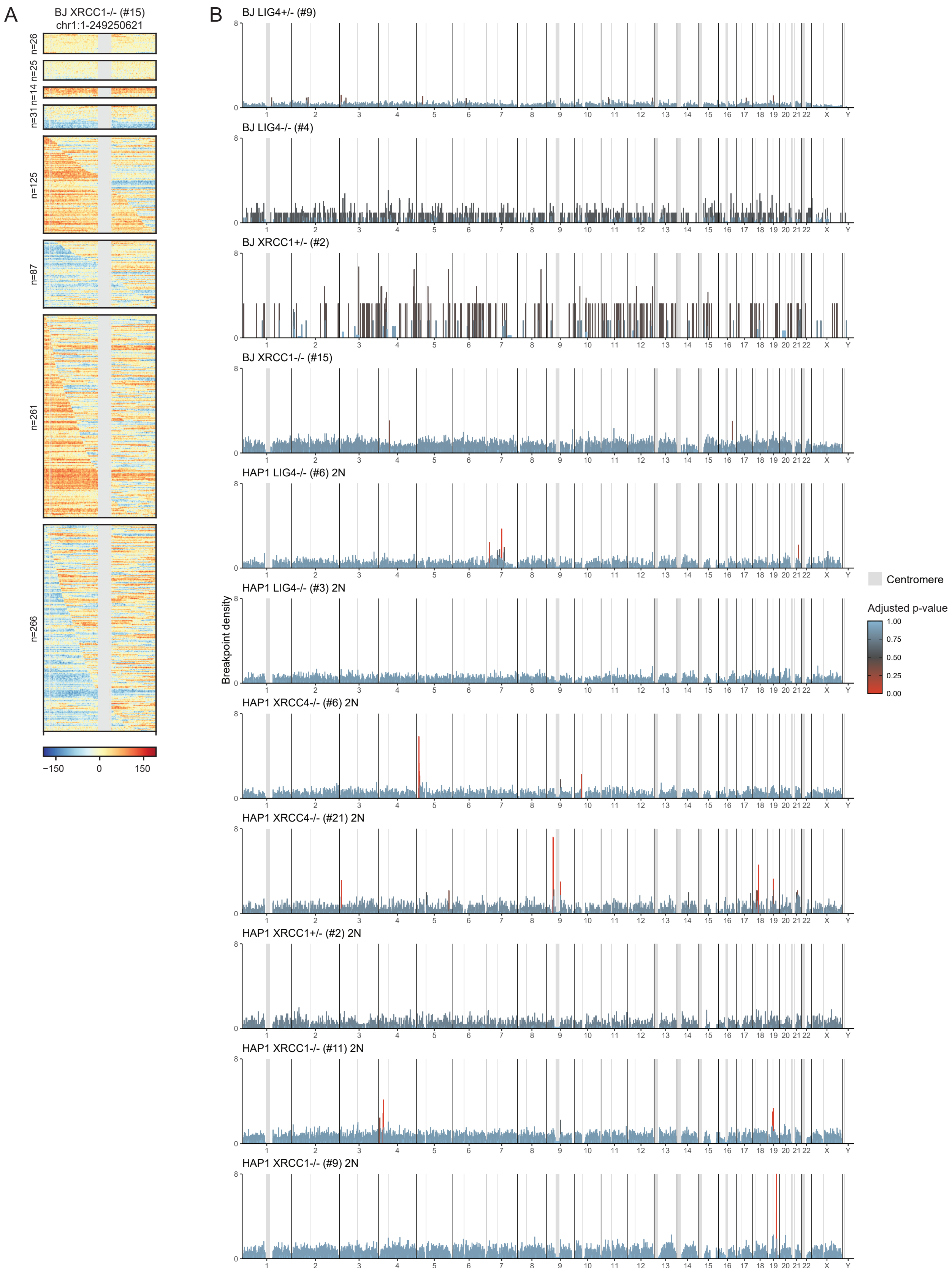
A
BJ XRCC1-/- (#15)
chr1:1-249250621
n=26
n=25
n=14
n=31
n=125
n=87
n=261
n=266
−150
0
150
B
BJ LIG4+/- (#9)
BJ LIG4-/- (#4)
BJ XRCC1+/- (#2)
BJ XRCC1-/- (#15)
HAP1 LIG4-/- (#6) 2N
HAP1 LIG4-/- (#3) 2N
HAP1 XRCC4-/- (#6) 2N
HAP1 XRCC4-/- (#21) 2N
HAP1 XRCC1+/- (#2) 2N
HAP1 XRCC1-/- (#11) 2N
HAP1 XRCC1-/- (#9) 2N
Breakpoint density
8
0
1 2 3 4 5 6 7 8 9 10 11 12 13 14 15 16 17 18 19 20 21 22 X Y
Centromere
Adjusted p-value
1.00
0.75
0.50
0.25
0.00

**Figure S3.** Genome-wide distribution of SCEs across DNA repair–deficient monoclonal knockout lines. **A.** Strand-states of chromosome 1 sorted by first break position across BJ-5ta XRCC1$^{-/-}$ cells. Each row represents a single cell with roughly 200 kb bin showing the normalized W-C read counts. Masked regions are in gray. Cells are divided into panels based on the number of breakpoints present, with top 4 panels having no breakpoints, next two having 1 breakpoint, and the last two having more than 1 breakpoint. **B.** Genome-wide SCE density for each knockout line. The distribution is plotted at 1Mb resolution, normalized by the number of bins each SCE overlaps with, and the total number of cells. Centromeres are highlighted in gray.

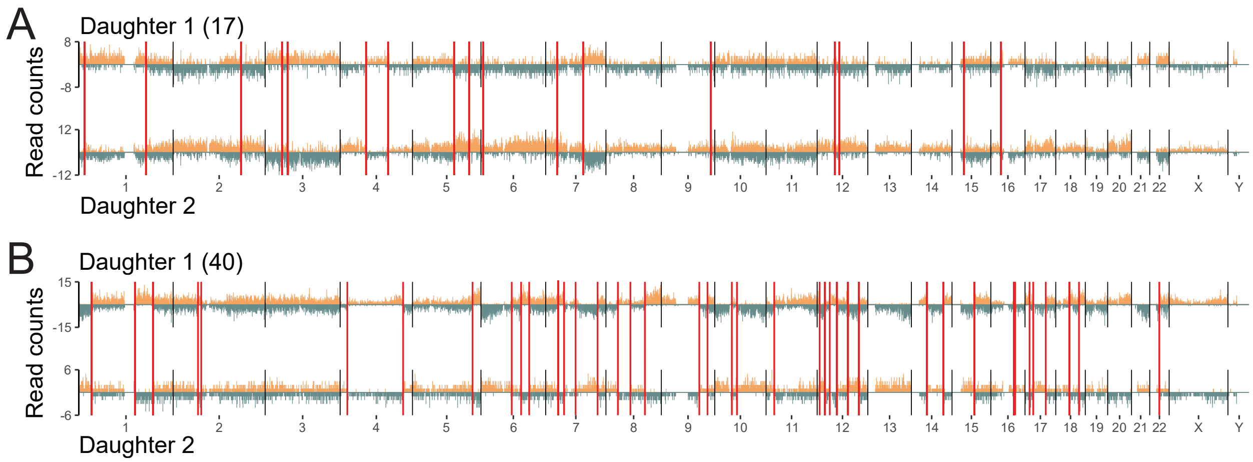


Figure S4. Additional examples of reciprocal daughter cells. Each vertical red line highlights a reciprocal SCE.
